# CLIC-dependent internalization of caveolin-1 to lysosomal vacuoles in response to osmotic regulation

**DOI:** 10.1101/2025.08.26.672461

**Authors:** Timothy H. Wong, Matheus F. Lima, Aditya Nagrath, Y. Lydia Li, Bharat Joshi, FuiBoon Kai, Ivan Robert Nabi

**Affiliations:** Department of Cellular & Physiological Sciences, Life Sciences Institute, University of British Columbia, Vancouver, BC, Canada V6T 1Z3; Department of Physiology and Pharmacology, Cumming School of Medicine, University of Calgary, Calgary, Alberta, Canada.; School of Biomedical Engineering, University of British Columbia, Vancouver, BC, Canada V6T 1Z3

**Keywords:** caveolae, caveolin-1, CLIC endocytosis, lysosome, osmotic shock

## Abstract

Originally thought to be a major endocytic portal, caveolae are now considered to function as a membrane buffer, whereby caveolae flattening protects the plasma membrane from rupture under mechanical stress, such as hypotonic shock. However, the fate of the caveolae coat protein caveolin-1 in response to hypotonic shock is not known. Here, we show that extended hypotonic shock induces ubiquitin-independent, CLIC-dependent endocytosis of caveolin-1 to large, intracellular, CD44-positive, pH-neutral lysosomal vacuoles negative for multivesicular body markers. Caveolin-1 recycles from these vacuoles to the plasma membrane upon return to isotonic conditions. Caveolin-1 internalizes not in response to initial cell expansion upon hypotonic shock but rather subsequent reduction of cell volume, and also in low-tension cells grown on reduced stiffness hydrogels. Upon hypertonic shock, caveolin-1 internalization occurs selectively in PC3 cells, lacking cavin-1, required for caveolae formation, and is inhibited upon cavin-1 reintroduction. CLIC endocytosis of non-caveolar caveolin-1 to neutral pH lysosomal vacuoles in response to osmotic regulation defines a novel recycling pathway enabling caveolae membrane buffering.

## Introduction

Caveolae were first identified by EM as smooth invaginations of the plasma membrane in the 1950’s and thought to be an endocytic portal, particularly in endothelial cells ^1-3^. However, the contribution of caveolae endocytosis to total endocytosis is relatively low and specific cargo for caveolae endocytosis have been difficult to identify, challenging the role of caveolae as a major endocytic portal ^3,4^. Caveolae were previously suggested to mediate the endocytosis of SV40 virus to a distinct Cav1-positive organelle, the caveosome, prior to viral delivery to the ER ^5^. Caveosomes were considered a distinct endocytic organelle as they were pH neutral and not labeled for early endosomal or lysosomal markers ^5^. However, the same authors subsequently identified the ubiquitin-dependent endocytosis of Cav1, showing that Cav1 ubiquitination promoted its endocytosis via multivesicular bodies to lysosomes for degradation ^6^. The caveosome was therefore considered to be an artifact of Cav1-GFP overexpression ^6,7^.

Indeed, a major function of caveolae is now considered to be not endocytosis but rather a membrane buffer, in which flattening of caveolae protects the plasma membrane from rupture under mechanical stress, such as hypotonic shock or cell stretching ^8-11^. In response to hypotonic shock, caveolae flatten, dissociate from the Cavin-1 adaptor required for caveolae formation, resulting in disassembly and release of dynamic Cav1 domains in the plasma membrane ^8,12-14^. Cav1 forms non-caveolar membrane structures or Cav1 scaffolds ^15^, including the disk-like 11-mer 8S complex recently characterized by cryoEM as well as larger 8S homo-oligomers ^16-18^. Dissociation of Cavin-1 from caveolae upon hypotonic shock is associated with increased dynamics of the released non-caveolar Cav1 scaffold domains as well as scaffold interaction with and inhibition of JAK-STAT signaling ^14,19^. However, the fate of disassembled Cav1 in response to hypotonic shock and how Cav1 can reform caveolae to protect the membrane against multiple cycles of stretch is not known.

Osmotic regulation is essential for maintaining cellular and tissue homeostasis, and in cancer, this balance is frequently disrupted due to alterations in membrane transport proteins, leading to dysregulated ion and water flux ^20,21^. Edema, which is characterized by elevated hydrostatic pressure and a hypo-osmotic microenvironment, often correlates with poor patient survival ^22,23^. Conversely, intratumoral hyperosmotic environments can trigger metabolic shifts that promote tumor progression ^24,25^.

Here, we show that extended osmotic shock results in the CLIC-dependent endocytosis of Cav1 to form large, intracellular, pH neutral, Cav1-positive lysosomal vacuoles that resemble the previously described caveosome. Cav1 internalization occurs independently of cavin-1, and therefore of caveolae, and of Cav1 ubiquitination; our data suggest that Cav1-positive lysosomal vacuoles are not an endosomal sorting station but rather a recycling lysosomal reservoir. Indeed, Cav1 endocytosis occurs not in response to cell expansion due to hypotonic shock but rather upon subsequent reduction of cell volume, consistent with previous reports of upregulated CLIC endocytosis in response to reduced cell volume ^26^. Cav1 scaffold endocytosis to Cav1-positive, pH neutral, lysosomal vacuoles defines a robust non-caveolar Cav1 recycling pathway to enable multiple rounds of caveolae buffering in response to osmotic regulation of cell volume.

## Results

Caveolae have been demonstrated to flatten in epithelial HeLa cells, mouse lung endothelial cells (MLECs), mouse embryonic fibroblasts (MEF) and mouse brain endothelial derived bEnd5 cells treated with acute (5-15 minutes) hypotonic shock, protecting the membrane from rupture in response to mechanical stress ^8,9,13^. Here, we used fixed cell TIRF to analyze the cell surface response of endogenous Cav1 to hypotonic shock over 60 minutes in MDA-MB-231 breast cancer cells. Cav1 surface expression progressively decreased over 60 minutes compared to cells imaged under isotonic conditions and recovered after 60 minutes upon addition of isotonic medium (Figure 1A). Using fixed cell STED imaging of Cav1, we observed Cav1 puncta at the cell membrane under isotonic conditions; in cells treated with hypotonic shock, Cav1 localized to large spherical vacuoles that appeared at 15 minutes of hypotonic shock and increased in abundance after 60 minutes (Figure 1B). These vacuoles were not visible by TIRF microscopy, suggesting that they are intracellular and do not correspond to the recently described doline ^27^. In the absence of cavin-1, Cav1 has been demonstrated to be endocytosed with CD44 via the CLIC/GEEC pathway ^28^. We therefore tested whether the hypotonic induced Cav1-positive vacuoles were positive for CD44 and the CLIC/GEEC cargo, β1 integrin, whose expression, signaling and endocytosis are regulated by Cav1 ^29-33^. While both CD44 and β1-integrin show a surface distribution under isotonic conditions, following 60 minutes of hypotonic shock, both CD44 and β1-integrin localize to Cav1 spherical vacuoles (Figure 1C). The average diameter of spherical vacuoles positive for Cav1, CD44, and β1-integrin was 0.97 µm and ranged from ∼0.3-3µm (Figure 1C). Dominant negative CDC42 has been shown to be a specific inhibitor of CLIC/GEEC endocytosis ^34,35^. Expression of wild-type CDC42 did not impact Cav1 endocytosis and CDC42-GFP associated with Cav1 and CD44 endocytic vacuoles. In contrast, dominant negative CDC42 mutant prevented the formation of Cav1 vacuoles, suggesting that these vacuoles are formed via CLIC endocytosis (Figure 1D and Supplemental Fig. 1).

**Figure 1.**
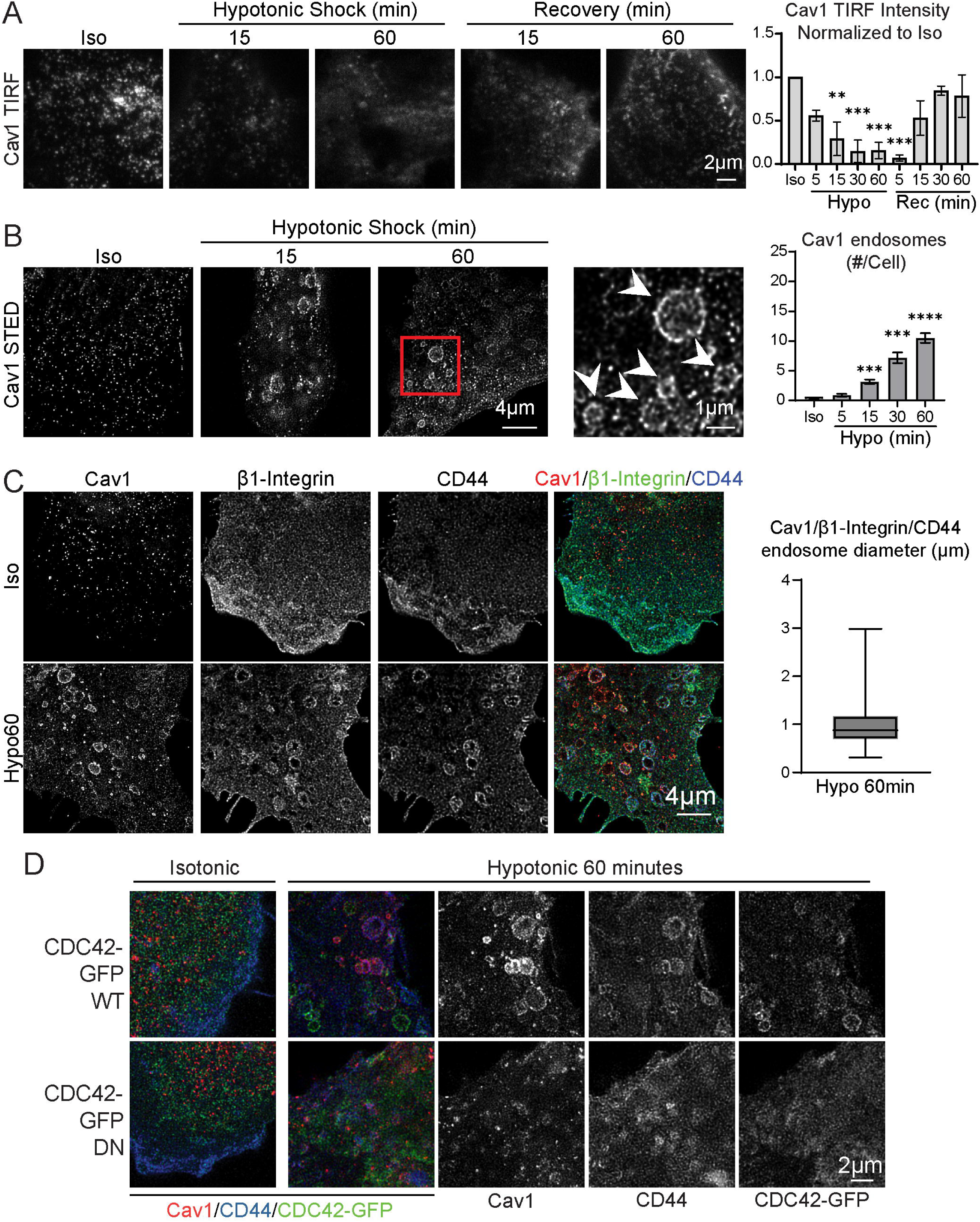
Hypotonic shock induces reversible internalization of Cav1 to vacuoles. A) MDA-MB-231 cells were incubated in isotonic media, in hypotonic media for 5, 15, 30 and 60 minutes or under Recovery conditions in which hypotonic media was replaced after 60 minutes with isotonic media for 5, 15, 30 or 60 minutes. Cells were labeled for Cav1, fixed and imaged by TIRF widefield imaging. Average Cav1 intensity per cell was quantified and normalized to isotonic control. Representative images of select conditions are shown. (n>30 whole cell images quantified for each condition from three independent experiments; ANOVA with Dunnett’s post-test comparing to isotonic control; **p < 0.01; ***p < 0.001; Scale bar: 2 µm). B) STED imaging of anti-Cav1 labeling in MDA-MB-231 cells incubated in isotonic media or in hypotonic media for 15 or 60 minutes. Arrow heads indicate Cav1 endocytic vacuoles. Cav1 endocytic vacuoles were counted per cell from whole cell images. (n>28 from three independent experiments for isotonic and hypotonic 15 minutes, four independent experiments for other conditions; (ANOVA with Dunnett post-test comparing to isotonic control; *p < 0.05; ****p < 0.0001; Scale bar: 4 µm; Inset scale bar: 1µm). C) MDA-MB-231 cells incubated with isotonic or hypotonic media for 60 minutes were fixed and labeled for Cav1, CD44 and β1-integrin (Scale bar: 4µm). The diameter of individual Cav1, CD44, and β1-integrin positive endosomes was measured from three independent experiments. D) MDA-MB-231 cells were transfected with wild-type (WT) CDC42-GFP or dominant negative (DN) CDC42-GFP, incubated with isotonic or hypotonic media for 60 minutes and then fixed and labeled for Cav1 and CD44.

To confirm that these spherical vacuoles are endocytic in nature, we performed an acid wash internalization assay using anti-CD44 and anti-β1 integrin antibodies (Figure 2A). Antibodies were added to hypotonic or isotonic media and at the indicated times, cells were acid-stripped to remove the surface antibody labelling and then fixed. As a control, antibodies were added for 60 minutes in isotonic media at 4°C to prevent endocytosis. Fluorescent labeling showed surface labeling of CD44 and β1 integrin without acid wash, that was completely removed by acid stripping at 4°C, demonstrating that the acid wash is removing surface labeling of CD44 and β1 integrin. Significantly increased internalization was detected for β1 integrin and CD44 antibodies at 15-, 30- and 60 minutes post hypotonic shock relative to internalization under isotonic conditions for 60 minutes (Fig. 2A; Supplemental Fig. S2A). Cav1 endocytic vacuoles are observed to be associated with internalized CD44 and β1 integrin (Fig. 2A, insets). Extent of overlap of a Cav1 mask with internalized CD44 and β1 integrin labelling increased over time in hypotonic medium and was significantly increased at 15-, 30- and 60 minutes post hypotonic shock relative to isotonic medium at 37°C (Fig. 2A). The CLIC inhibitor 7-Ketocholesterol (7-KC) ^36,37^ prevented internalization of CD44 antibodies as well as uptake of Cav1, as determined by Cav1 overlap with internalized CD44, after 60 minutes of hypotonic shock (Fig. 2B; Supplemental Fig. S2B). Consistently, Cav1/CD44 positive vacuoles are positive for the CLIC endocytosis reporter, GFP-GPI ^38,39^ and negative for the clathrin endocytosis marker transferrin receptor (Supplemental Fig. S3A). The GLUT1 glucose transporter, which undergoes both clathrin and non-clathrin dependent Arf6 dependent endocytosis ^40-42^, associates with CD44 after 60 minutes of hypotonic shock (Supplemental Fig. S3A). 3D structured illumination (SIM) microscopy of acid washed cells showed that after 2 minutes of hypotonic shock anti-CD44 antibodies colocalized with Cav1 in short, tubular structures corresponding to CLIC endocytic tubules. XZ scans show that CLIC tubules are present around the cell surface; after 60 minutes of hypotonic shock, spherical Cav1 and CD44-positive vacuoles are observed throughout the cell (Figure 2C). In response to hypotonic shock, Cav1 is therefore endocytosed via the CLIC/GEEC endocytic pathway.

**Figure 2.**
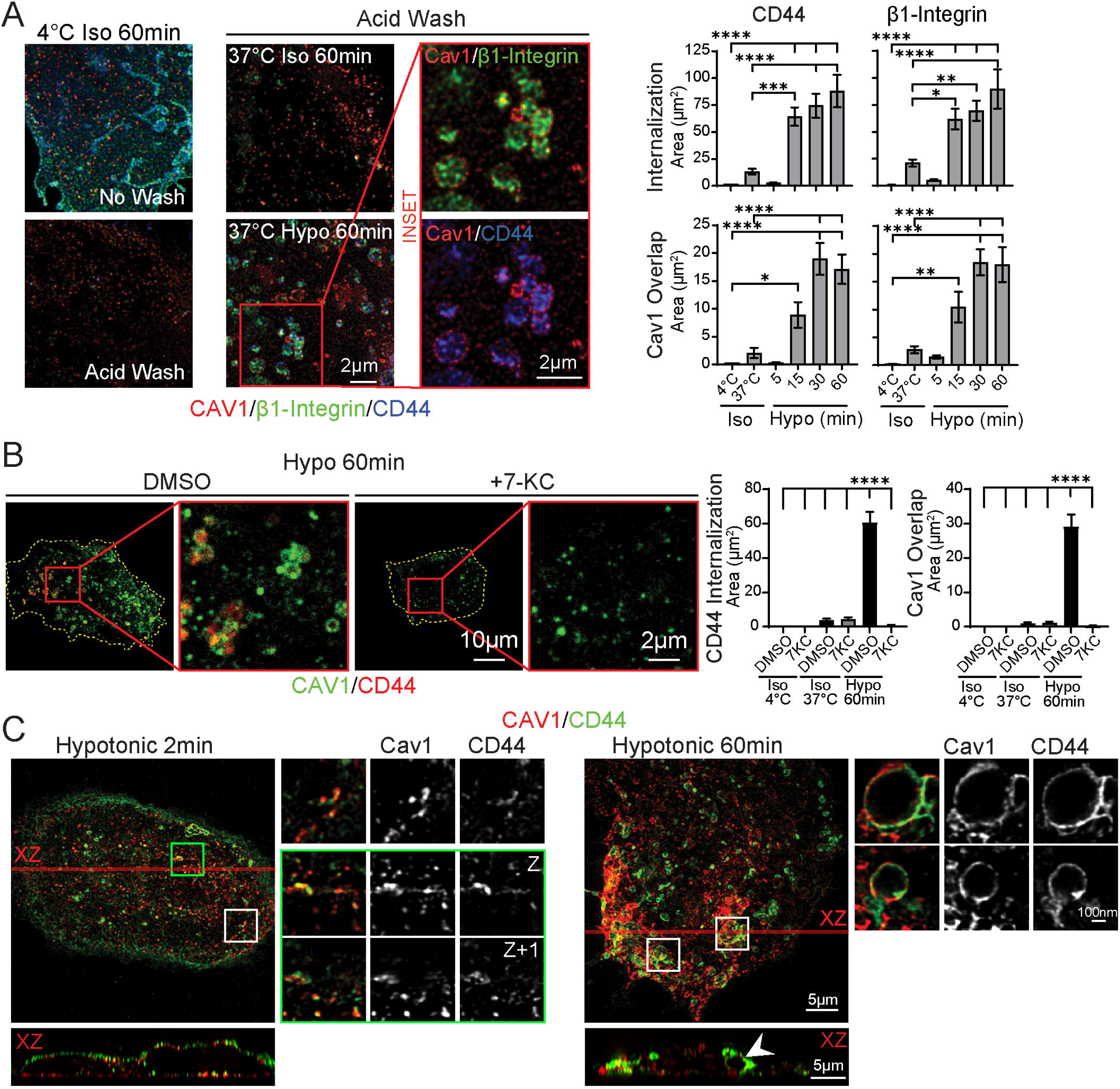
Cav1 is endocytosed via the CLIC pathway in response to hypotonic shock. A) MDA-MB-231 cells were incubated with anti-CD44 and anti-β1 integrin antibodies in isotonic media for 60 minutes at 4°C or 37°C or in hypotonic media for 5, 15, 30 or 60 minutes at 37°C. As indicated, an acid wash treatment was performed at 4°C to remove surface antibodies and cells were fixed and labeled with secondary antibodies to anti-CD44 and anti-β1 integrin as well as for endogenous Cav1. (Quantified from whole cell images, n≥30 cells per condition from three independent experiments; ANOVA with Dunnett post-test comparing each condition to isotonic 4°C control; *p < 0.05; ***p < 0.001; ****p < 0.0001; Scale bars: 2 µm). B) Confocal imaging of MDA-MB-231 cells pre-treated with the CLIC inhibitor 7-Ketocholesterol (7-KC) and then incubated with anti-CD44 antibodies and acid washed. Internalized anti-CD44 and overlap with Cav1 were quantified (n = 30 cells per condition from three independent experiments; ANOVA with a Tukey post-test; ****p < 0.0001) C) 3D SIM imaging of MDA-MB-231 cells incubated with anti-CD44 antibodies in hypotonic media for 2 or 60 minutes and then acid washed and labeled for Cav1. Maximum projections of the Z stack are shown with insets (boxes) taken from individual z-slices. The green ROI and inset depict CLIC tubules from two adjacent z-slices. XZ sections were generated along the transparent red line and show internalized CD44 (green) and its association with Cav1. Arrowheads indicate the Cav1 vacuole depicted in the XZ sections of the hypotonic shock 60minutes ROI (120 nm step size; Scale bar: 500nm; Inset scale bar: 100nm).

Ubiquitin-dependent trafficking of Cav1 to lysosomes via multivesicular bodies (MVBs) has been reported ^6^. We therefore expressed LAMP1-GFP and found that upon hypotonic shock more than 50% of Cav1 endocytic vacuoles colocalized with LAMP1-GFP (Figure 3A). To determine whether ubiquitination is required for Cav1 endocytic vacuole formation, we expressed in Cav1 knockout MDA-MB-231 cells ^43^, wild-type Cav1-HA or the Cav1-K*R-HA mutant in which all lysine residues are mutated to arginine, that was previously shown to prevent Cav1 ubiquitination and internalization to lysosomes ^6^. In response to hypotonic shock, the Cav1-K*R-HA ubiquitin mutant localized to Cav1-positive endocytic vacuoles to the same extent as wild-type Cav1-HA, indicating that Cav1 ubiquitination is not required for osmotic regulated Cav1 endocytosis (Figure 3B and Supplemental Fig. S4). Consistently, Cav1/CD44 positive vacuoles are labelled for endogenous LAMP2 and negative for the MVB markers CD63 and LBPA, further supporting that Cav1/CD44 vacuoles are not MVBs (Supplemental Figure S3B).

**Figure 3.**
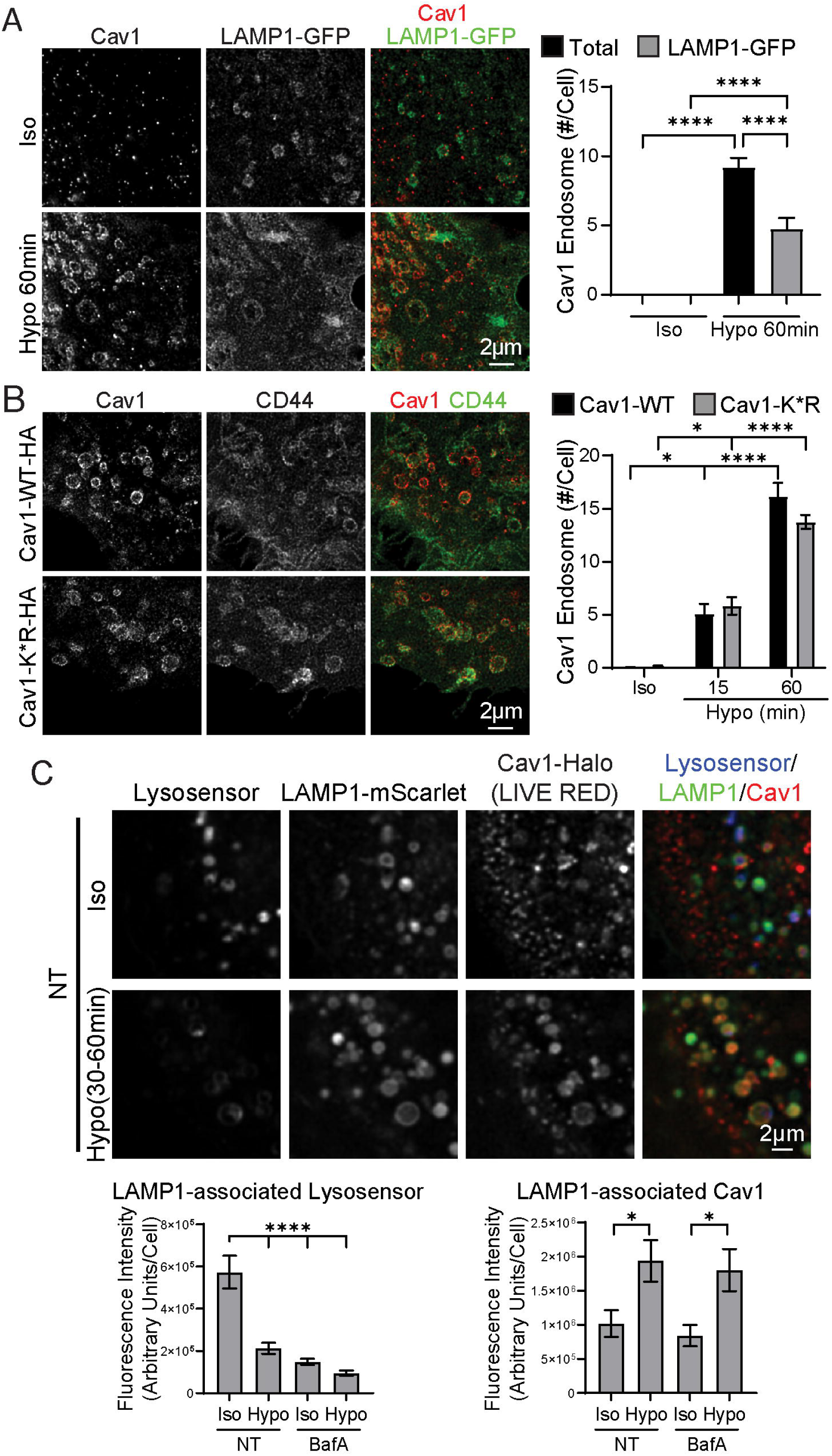
Ubiquitin-independent endocytosis of Cav1 to pH neutral lysosomes. A) MDA-MB-231 cells were transfected with LAMP1-GFP, incubated with isotonic or hypotonic media for 60 minutes and then fixed and labeled for Cav1. From 2D STED images, the total number of Cav1 endocytic vacuoles and those colocalizing with LAMP1-GFP were counted per cell (n>30 cells for each condition from three independent experiments; ANOVA with Tukey’s post-test; ****p < 0.0001; Scale bar: 2 µm). B) Cav1 CRISPR knockout MDA-MB-231 cells were transfected with Cav1-WT-HA or with the Cav1-K*R-HA lysine mutant, incubated with isotonic for 60 minutes or hypotonic media for 15 or 60 minutes and then fixed and labeled for Cav1 and CD44. The number of Cav1 endocytic vacuoles was counted per cell (n>29 cells from three independent experiments; ANOVA with Tukey post-test comparing WT to K*R for the same condition and for hypotonic relative to isotonic; Scale bar: 2µm). C) MDA-MB-231 cells were transfected with LAMP1-mScarlet and Cav1-Halo, pre-treated with 100 nM BafA for 2 hours where indicated, labeled with Halo-LIVE RED for 30 minutes and then incubated for 30 minutes with 1 μM LysoSensor green and LIVE RED Halotag ligand. Cells were then placed in isotonic or hypotonic imaging media and multiple cells imaged over a period of 30 minutes in isotonic DMEM or after 30 minutes of hypotonic shock. LysoSensor and Cav1 intensity within LAMP1-positive lysosomes was quantified. (n>38 cells from three independent experiments; ANOVA with Tukey’s post-test; *p < 0.05; ****p < 0.0001; Scale bar: 2 µm).

To determine whether Cav1-positive lysosomal vacuoles are acidic, we performed live cell experiments of MDA-MB-231 Cav1 knockout cells ^44^ expressing Halo-tagged Cav1 labeled with Halo ligand LIVE RED, and LAMP1-mScarlet and incubated the cells with the acid-dependent lysosomal reporter LysoSensor green. Under isotonic conditions, LAMP1-mScarlet lysosomes are strongly labeled for both LAMP1 and LysoSensor and poorly labeled for Cav1-Halo. However, under hypotonic shock, LAMP1-mScarlet lysosomes are strongly labeled for Cav1-Halo and poorly labeled for LysoSensor (Figure 3C). Similarly, upon treatment with bafilomycin A1 (BafA), a V-ATPase inhibitor that neutralizes endolysosomal pH, cells under isotonic conditions contain LAMP-1-positive lysosomes poorly labelled for LysoSensor. Further, BafA inhibition of lysosome acidification did not prevent the formation of Cav1 lysosomal vacuoles. Taken together, these results indicate that in response to hypotonic shock Cav1 is trafficked into pH-neutral lysosomal vacuoles independently of Cav1 ubiquitination.

Assessment of cell volume changes using quantitative phase imaging microscopy showed that cell volume spiked within five minutes after hypotonic shock and then decreased below the initial isotonic cell volume, plateauing at a reduced cell volume after about 15 minutes of hypotonic shock (Figure 4A). These results suggest that Cav1 endocytosis does not occur during initial cell swelling upon hypotonic shock but rather during the subsequent decrease in cell volume. To investigate whether Cav1 endocytic vacuoles form in response to decreasing cell volume, we applied both hypertonic and hypotonic shock to MDA-MB-231 cells as well as PC3 prostate cancer cells, that lack cavin-1 and do not form caveolae ^45^. MDA-MB-231 cells accumulated spherical Cav1-positive lysosomal vacuoles over 15-60 minutes post-hypotonic shock as described previously, but not upon hypertonic shock. In PC3 cells, Cav1-positive lysosomes formed at 5 and 15 minutes post-hypotonic shock and, in contrast to MDA-MB-231 cells, Cav1-positive lysosomal vacuoles accumulated over time in PC3 cells under hypertonic shock (Figure 4B; Supplemental Fig. S5A). To test whether cavin-1 expression is limiting Cav1 endocytosis in response to hypertonic shock, we expressed cavin-1-GFP in PC3 cells. Under hypertonic shock, cells expressing cavin-1-GFP no longer formed Cav1 lysosomal vacuoles, whereas under hypotonic shock, Cav1 still trafficked to lysosomal vacuoles (Figure 4C; Supplemental Fig. S5B). TIRF microscopy showed that in both MDA-MB-231 cells and cavin-1-expressing PC3 cells, cavin-1 surface expression and association with Cav1 decrease at 5 minutes post-hypotonic shock, whereas Cav1 is significantly lost from the membrane only at 15 minutes hypotonic shock (Supplemental Fig. S6). Hypotonic shock induced Cav1-cavin-1 dissociation is therefore required for CLIC-dependent Cav1 endocytosis in response to cell shrinkage.

**Figure 4.**
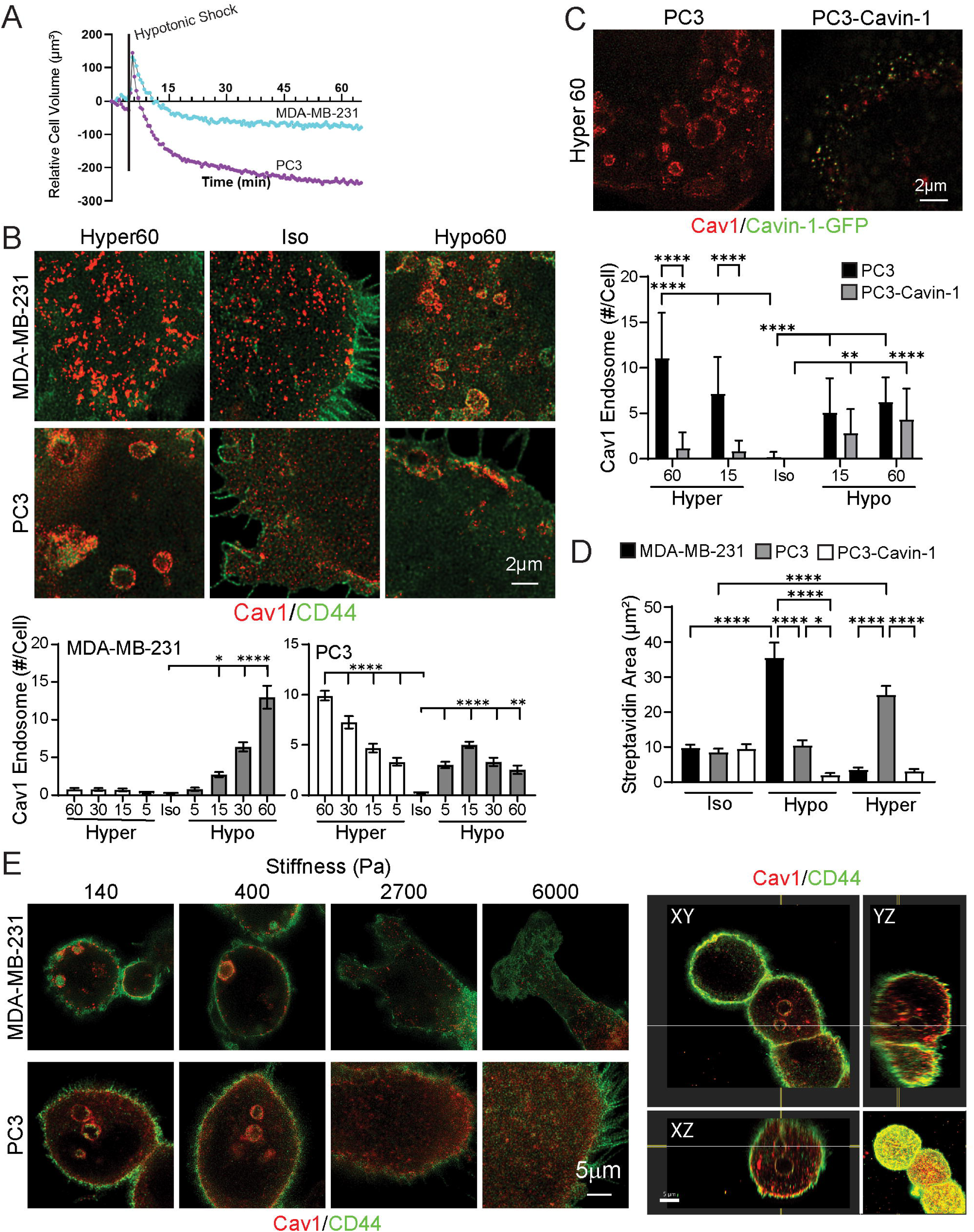
Cavin1-independent Cav1 endocytosis is associated with reduced cell volume. A) Cell volume of MDA-MB-231 and PC3 cells switched from isotonic to hypotonic media was measured by live phase imaging. The black line indicates addition of hypotonic media (three independent experiments, each including 2-3 technical replicates per sample with each technical replicate consisting of 9 ROIs). B) MDA-MB-231 and PC3 cells were incubated in isotonic media for 60 minutes, or hypotonic or hypertonic media for 5, 15, 30 and 60 minutes. Cells were fixed and labeled for Cav1 and CD44 and Cav1 endocytic vacuoles counted per cell from whole cell images. Representative images for select time points are shown. (n>28 cells from three independent experiments; ANOVA with Dunnett’s post-test comparing each condition to isotonic control; *p < 0.05; **p < 0.01; ****p < 0.0001; Scale bar: 2 µm). C) PC3 cells and PC3 cells transfected with Cavin-1-GFP were incubated in isotonic, hypotonic and hypertonic shock media for 60 minutes and then fixed and labeled for Cav1. Cavin-1-GFP transfected cells were identified in the GFP channel and Cav1 endocytic vacuoles counted per cell. (n>28 cells from three independent experiments; ANOVA with Tukey’s post-test and comparing PC3 to PC3-Cavin-1 and each condition with the respective isotonic control; **p < 0.01; ***p < 0.001; ****p < 0.0001; Scale bar: 2 µm). D) Plasma membrane proteins of MDA-MB-231, PC3 cells and PC3 cells transfected with Cavin1-GFP were biotinylated with Sulfo-NHS-SS-biotin at 4°C and after incubation in isotonic, hypotonic and hypertonic shock media for 60 minutes at 3°7C surface biotin was removed by glutathione (GSH) wash. Cells were labeled with Alexa568-conjugated streptavidin and internalized biotin quantified (n = 30 cells per condition from three independent experiments; ANOVA with a Tukey post-test comparing isotonic conditions to hypo and hypertonic shock as well as comparisons between cell lines for each respective osmotic condition; *p < 0.05; ****p < 0.0001). E) MDA-MB-231 and PC3 cells were seeded for 24h on fibronectin conjugated PA hydrogel substrates with the indicated stiffnesses before fixation and labelling for Cav1 and CD44. Representative images from 3 independent experiments are shown. XY, XZ and YZ cross-section views of a 3D STED image stack of PC3 cells grown on 140Pa stiffness hydrogel show an intracellular Cav1 and CD44 vacuole.

To test whether osmotic shock similarly impacted global plasma membrane endocytosis, cells were surface labeled with Sulfo-N-Hydroxysulfosuccinimide-SS-biotin (Sulfo-NHS-SS-Biotin) at 4°C and, treated with the reducing agent l-glutathione reduced (GSH) to remove surface-bound biotin after 60 minutes incubation in isotonic or osmotic shock media at 37°C. Plasma membrane protein endocytosis measured by internalized biotin, increased after 60 minutes upon hypotonic shock but not under hypertonic shock in MDA-MB-231 cells (Fig. 4D; Supplemental Fig. S7). In PC3 cells, plasma membrane protein endocytosis was induced upon hypertonic shock and inhibited by expression of cavin-1 (Fig. 4D).

To determine whether other conditions that alter cell volume or shape similarly promote Cav1 vacuole formation, cells were seeded on fibronectin-conjugated polyacrylamide (PA) hydrogels with defined elastic moduli spanning a range of physiologically relevant stiffnesses (Fig. 4E and Supplemental Fig. S8). In both MDA-MB-231 and PC3 cells, Cav1- and CD44-positive vacuoles were readily observed on compliant PA hydrogels (140 and 400 Pa). In contrast, cells cultured on stiffer hydrogels (2.7 and 6 kPa) exhibited a more spread morphology and did not form vacuoles. As observed with acute osmotic shock, chronic exposure to compliant ECM substrates promotes Cav1 endocytosis into CD44-positive vacuoles.

## Discussion

We show here that in response to extended hypotonic shock, Cav1 is robustly and reversibly internalized via CLIC endocytosis to neutral pH lysosomal vacuoles in a caveolae-independent manner. Loss of cavin-1 and caveolae disassembly increase Cav1 lateral mobility and rate of degradation ^6,45,46^. However, we observe that Cav1 lysosomal vacuoles are negative for the acidic pH indicator Lysosensor and upon return to isotonic conditions, Cav1 efficiently recycles to the plasma membrane. Furthermore, a lysine-free Cav1 mutant, unable to be ubiquitinated, is efficiently internalized in response to hypotonicity; osmotically regulated Cav1 endocytosis is therefore distinct from ubiquitin-dependent endocytosis of Cav1 to lysosomes for degradation ^6^. Tracking cell volume in response to hypotonic shock demonstrates that Cav1 endocytosis and endocytic vacuole formation are occurring as cells acclimatize to hypotonic shock and decreased cell volume. Consistently, other studies measuring cell volume in response to hypotonic shock show that ion channel compensation decreases cell volume in response to the initial volume spike caused by hypotonic shock ^47,48^. Similarly, Cav1 vacuole formation is also observed in cells seeded on compliant ECM. Indeed, cell growth on soft substrates, associated with reduced cell spreading and volume, has been reported to be associated with Cav1 endocytosis ^49,50^.

Our data define a Cav1 recycling pathway in response to hypoosmotic shock: 1. caveolae flatten to buffer membrane stretch and release cavin-1 to form Cav1 scaffolds; 2. scaffolds then internalize via CLIC endocytosis to pH neutral lysosomal vacuoles; and 3. Cav1 recycles to the plasma membrane to enable reformation of caveolae. Cav1 has been shown to be internalized via the CLIC pathway in the absence of cavin-1, and indeed cavin-1 shown to inhibit its endocytosis ^28^. Consistently, in PC3 cells lacking cavin-1, Cav1 endocytosis occurs in response to hyperosmotic shock and is prevented by reintroduction of cavin-1. The CLIC pathway has a high capacity for endocytosis, responds to and regulates membrane tension and is a major contributor to fluid-phase uptake in response to mechanical stress ^26,51^. Interestingly, vinculin regulates fluid uptake via CLIC/GEEC during recovery following osmotic shock ^26^, Cav1 modulates vinculin focal adhesion tension ^52^ and Cav1 inhibits CLIC endocytosis ^28^. Cav1 therefore sits at the intersection of two mechanosensitive mechanisms, caveolae flattening and CLIC/GEEC endocytosis, both implicated in the cellular response to mechanical and osmotic stress. CLIC-dependent, non-caveolar endocytosis of Cav1 to neutral pH lysosomal vacuoles in response to osmotic changes represents a robust recycling pathway for Cav1 in response to altered membrane tension that may enable repeated caveolae membrane buffering.

While the Cav1-positive neutral pH lysosomal vacuoles reported here resemble the previously described “caveosome” ^5^, they are not an endocytic sorting compartment but rather an intracellular recycling compartment for cellular management of cell volume changes in response to changes in osmotic pressure. Indeed, LAMP-1-positive and Lysotracker-negative vacuoles were recently reported to form 30 minutes following hypotonic shock, accumulating over 6 hours ^53^. Upon hypotonic shock, lysosomes can act as intracellular water storage vacuoles regulated by the volume-regulated anion channel LLRC8A, contributing to cell volume decreases and cell survival ^46,54^. Rapid caveolae endocytosis has been reported to repair plasma membrane wounds in association with Ca^++^-dependent exocytosis of lysosomes; loss of Cav1 impairs wound repair and is associated with enhanced Endophilin A2 endocytosis ^55-57^. How this membrane repair pathway of Cav1 endocytosis relates to that reported here in response to osmotic shock is not clear, however our data argue that Cav1 endocytosis in response to osmotic shock is scaffold, and not caveolae, mediated. Of interest, downregulation of both Cav1 and cavin-1 were shown to increase the expression of matrix proteases and glioblastoma invasion in response to increased hydrostatic pressure ^58^. Whether Cav1 impacts the dynamics or function of this organelle or is simply a bystander cargo internalized upon induction of CLIC endocytosis upon membrane relaxation for recycling to reform membrane caveolae remains to be determined.

## Method Details

### Plasmid generation

In order to generate dually tagged Halo and Myc human Cav-1, Cav1-myc and Halo Tag containing pST018 (a gift from Sheila Teves, UBC) were used as template DNAs and PCR (Platinum Taq, Invitrogen) amplified following overlapping extension method using a set of forward and reverse primers (5’GGA AGC TTA GCA TGT CTG GGG GCA AAT AC3’ 5’CGG CCG GTC GCC ACC ATG GCA GAA ATC GGT3’; 5’ACC GAT TTC TGC CAT GGT GGC GAC CGG CCG3’, 5’GAA TTC TTA CGT ACG CTT GTC GTC ATC GTC TTT GTA GTC3’). The PCR products were TA cloned and plasmids sequence verified to check fusion of Halo tag with Cav1-Myc. The correct sequence containing Cav1-Myc-Halo fragment was released from the TA-clone using EcoR1 and re-cloned into pcDNA3 at the EcoR1 site. Positive clones were sequence verified for the correct orientation and used for the transfection and subsequent experiments.

### Cell culture, transfection, osmotic shock and drug treatments

Generation and characterization of the cloned MDA-MB-231 and PC3 cell lines and the Cav1 knockout cell lines used in this study were as described previously ^44,59^. All cell lines were tested regularly for mycoplasma infection by PCR (ABM, Richmond, BC, Canada). Cell lines were grown in Gibco™ RPMI-1640 medium (CAT#21870076) that was supplemented using 10% fetal bovine serum (FBS) and 2 mM L-glutamine (ThermoFisher). MDA-MB-231 cells were passaged using 0.05% Trypsin-EDTA (ThermoFisher) and 0.25% for PC3 cell lines and used for a maximum of ten passages. For plasmid transfections, cells were seeded for 24 h and then transfected overnight using Lipofectamine 2000 (Life Technologies, Thermo Fisher Scientific) as described by the manufacturer’s protocol. For live cell confocal imaging, cells were seeded for 24 h on an 8-well #1.5H glass bottom slide (Ibidi). Transfection was performed overnight as described above.

Cells grown on #1.5H 18mm square coverslips (Paul Marienfeld, Lauda-Königshofen, Germany). were treated with osmotic shock after being seeded for 48 h. For hypoosmotic shock (30 mOsm), complete media (300 mOsm) was diluted 1:10 with MQ water and cells were treated for the time duration as described at 37°C, 5%CO2. For hyperosmotic shock (500 mOsm), 2M NaCl solution was diluted 1:10 with complete media. Where indicated, before imaging, cells were pre-treated with 100nM BafA (Sigma Aldrich 19-148) for a total of 2 h and kept in media containing BafA for subsequent incubations and for imaging. Cells were then incubated for 30 minutes with 1uM LysoSensor green (ThermoFisher L7535) and LIVE RED Halotag ligand (Abberior LVRED-0147-30NMOL) as indicated before being washed 3x with HEPES buffered DMEM without phenol red (ThermoFisher 21063-029). Multiple cells were imaged over a period of 30 minutes in isotonic DMEM or after 30 minutes of osmotic shock for live cell imaging.

### Acid-wash internalization assay

Internalization assays were adapted from Chaudhary *et al.* ^28^. Cells were grown on coverslips in 6 well plates before being transferred to a 150mm dish lined with parafilm. For Cells were treated with a 250 µl solution of isotonic or hypotonic media containing 10 µg/ml anti-CD44 (Novus 22530) and anti-β1 integrin antibody and incubated for the indicated times at 37°C, or at 4°C as indicated. To remove uninternalized and surface bound antibodies, acid stripping was performed using 2x30 second washes in 0.5 M glycine at pH 2.2. Cells were then fixed with 4% paraformaldehyde for 15 minutes and labeled with anti-Cav1 primary and secondary antibodies.

For 7-Ketocholesterol (7-KC) experiments, MDA-MB-231 cells were treated for 24 hours with 30 μM 7-KC with 0.1% DMSO used as the vehicle control. Cells were then incubated with 1.6 μg/mL mouse anti-CD44 (Cell Signaling 156-3C11) antibody for 60 minutes in either isotonic or hypotonic media. Acid stripping, cell fixation and labelling for Cav1 and anti-CD44 performed as described above.

### Polyacrylamide (PA) hydrogel preparation

PA gel substrates with defined elastic moduli (140 Pa: 3% acrylamide, 0.04% bis-acrylamide; 400 Pa: 3% acrylamide, 0.05% bis-acrylamide; 2.7 kPa: 7.5% acrylamide, 0.035% bis-acrylamide; 6 kPa: 7.5% acrylamide, 0.07% bis-acrylamide) were synthesized as previously described, with minor modifications ^60^. Fibronectin (20 μg/mL) was covalently conjugated to the PA substrates using 0.01% bis-acrylamide, 0.025% Irgacure 2959, 0.002% di(trimethylolpropane) tetracrylate (Sigma-Aldrich), and 0.01% N-succinimidyl acrylamidohexanoic acid. Following gel functionalization, gels were extensively rinsed with sterile buffer and equilibrated in complete culture medium at 37 °C prior to seeding 10,000 cells.

### Fluorescence labeling and imaging

Cells were fixed with 4% paraformaldehyde (Electron Microscopy Sciences) and permeabilized using 0.2% Triton X-100 (Sigma) diluted in PBS supplemented with 1 mM MgCl_2_ and 0.1 mM CaCl_2_ (PBS-CM). Blocking was performed for 1 h with 10% goat serum (ThermoFisher) and 1% bovine serum albumin (Sigma). Coverslips were incubated with primary antibodies (diluted in antibody buffer containing 10% goat serum, 1% BSA, and 0.05% Triton X-100 in saline-sodium citrate buffer) overnight at 4°C and then washed with antibody wash buffer containing saline-sodium citrate buffer and 0.05% Triton X-100. Fluorescent secondary antibodies were incubated for 1h at room temperature and washed with antibody wash buffer before being mounted with ProLong™ diamond antifade mountant (ThermoFisher). STED imaging was carried out using a Leica SP8 microscope with a 100x/1.4 Oil HC PL APO CS2 STED White objective. Live and fixed cell confocal imaging was performed using a Leica Stellaris 5 microscope using a 63x objective (HC PL APO 63x/1.40, oil immersion). STED and live cell confocal images were deconvolved using Huygens Professional software (Scientific Volume Imaging). TIRF imaging was performed using a Leica SR GSD 3D system using a 160 × objective lens (HC PL APO 160x/1.43, oil immersion) with TIRF illumination. 3D SIM imaging was performed with a ZEISS Lattice SIM 3 using a 40x objective lens (Plan-Apochromat 40x/1.4 Oil DIC (UV) VIS-IR M27).

### NHS-SS-Biotin Surface Labeling and Uptake

Cells were washed twice in ice-cold 1X PBS-CM followed by incubation with EZ-Link™ Sulfo-NHS-SS-Biotin (ThermoFisher, Cat N:21331) at 0.25 mg/ml for 15 minutes at 4°C with gentle shaking. Cells were washed twice in TBS (pH 7.4) followed by 2 washes in 100 mM glycine in 1X PBS-CM and then incubated with isotonic, hypotonic and hypertonic media for 1h at 37°C. Then, cells were washed twice in L-glutathione buffer (75 mM NaCl, 75 mM NaOH, 10 mM EDTA, 50 mM l-glutathione reduced (GSH) (Sigma Aldrich G6013) for 5 minutes at 4°C, with gentle shaking, to remove cell surface biotin and then washed 3 times in 50 μg/ml of iodoacetamide in PBS-CM to quench the GSH. Cells were then fixed, permeabilized and blocked as described above. Biotin was labelled with Alexa568-conjugated streptavidin diluted in antibody buffer and then washed and mounted as described above.

### Volume Phase Imaging

Cells were seeded for 48h on µ-Slide VI 0.5 Glass Bottom (1.5H) from Ibidi (80607). A gravity perfusion system was set up by adapting an Ibidi tubing set (10831) and perfusion set (10963). Briefly, the reservoirs from the Ibidi perfusion set were used to store isotonic and hypotonic solutions and configured with a Y tube fitting to combine into a single tube and connect with the µ-Slide. Tubing from each reservoir was clamped to control the release of the solutions. The Ibidi tubing set was used as the waste line to connect the slide to a collection Falcon tube. Imaging was performed with a HoloMonitor (PHI) quantitative phase imaging microscope placed inside a cell culture incubator culture incubator at 37°C and 5% CO_2_. Cells were imaged for 5 minutes under isotonic media and was exchanged with the prepared hypoosmotic media. 8-10 regions of interest (ROIs) were selected, and images were acquired every 30 seconds. Cells were segmented, tracked, and quantified at each timepoint using the HoloMonitor software package. A single cell tracking module was applied and 7-10 cells were selected from each ROI for quantification. Cells were selected based on the software being able to sufficiently isolate the single cell by segmentation and track the same cell for the entire duration of the experiment.

## Quantification and Statistical Analysis

Quantification of TIRF Cav1 fluorescence was performed with ImageJ/FIJI by measuring RawIntDen of the Cav1 channel for individual whole cells and normalizing the values to the isotonic control. Quantification of Cav1-positive endocytic vacuoles was done by manual scoring. Quantification of internalization assays and area overlap was performed using the ImageJ plugin JACoP ^61^. To quantify LysoSensor and Cav1 intensity within LAMP1-positive lysosomes, ImageJ/FIJI was used to generate a mask from the LAMP1-mScarlet channel. The mask was used to quantify the fluorescence intensity of Cav1 and Lysosensor channels within the mask for each cell. Graphs and statistical analyses were performed using GraphPad Prism (GraphPad Software, San Diego, CA).

Data are presented as mean ± SEM. The statistical test used, definition of significance, and number of biological replicates are reported in the figure legends. n denotes the number of cells analyzed from at least three independent biological replicates. Group comparisons were performed using one-way ANOVA with Dunnett’s or Tukey post-test as indicated in the figure legends. p-values <0.05 were considered statistically significant, with significance annotated as *p < 0.05, **p < 0.01, ***p < 0.001, ****p < 0.0001.

## Acknowledgements

This study was supported by a grant from the Canadian Institutes of Health Research (IRN: PJT-175112). We thank Christophe Lamaze for insightful comments and suggestions.

## Notes

### Competing Interest Statement

The authors have declared no competing interest.

### Summary of Updates

Key additions to the revised manuscript: Further validation of the CLIC pathway: We show that the CLIC inhibitor 7-Ketocholesterol prevents CD44 uptake and Cav1 endocytosis in response to osmotic shock (Fig. 2B and Supplemental Fig. S2) 3D SIM imaging: We include 3D SIM imaging localizing Cav1 to CLIC tubules during the early stage of hypotonic shock and representations of Cav1/CD44 vacuoles (Fig. 2C). Biotin surface labeling and uptake experiment: Using Sulfo-NHS-SS-biotin uptake, we show that global plasma membrane endocytosis follows Cav1 endocytosis to vacuoles upon osmotic shock (Fig. 4D and Supplemental Fig. S7). Physiological relevance: We expanded our study to include fibronectin-conjugated hydrogels of varying physiologically relevant stiffnesses, confirming that Cav1/CD44 vacuole formation is a general response to mechanical stimuli beyond osmotic shock (Fig. 4E and Supplemental Fig. S8). Other markers: We show that osmotic shock induced Cav1/CD44-positive vacuoles are positive for GPI-GFP and endogenous GLUT1, and negative for the clathrin-mediated endocytosis marker, transferrin Receptor (Supplemental Fig. S3A). Cavin-1 dissociation upon hypotonic shock: Supplemental Figure S6 now includes imaging data from both MDA-MB-231 and Cavin-1 expressing PC3 cell lines, demonstrating the dissociation of Cavin-1 from Cav1 upon hypotonic shock. Reorganized the manuscript to improve clarity and accommodate the new experiments: We include quantification of CDC42-GFP expression levels in Supplemental Fig. S1. To better organize the figures, we moved the original Figure 2B to Figure 1C, with the corresponding quantification to Supplemental Fig. S1. To accommodate new data in Figure 2, the "hypo15" image (formerly Fig. 2A) has been moved to Supplemental Fig. S2A. Figure 4 was reorganized for better spacing; representative "hypo15" and "hyper15" images (formerly 4B) and the PC3 and Isotonic images (formerly 4C) have been moved to Supplemental Fig. S5A and S5B, respectively. Previous and new data showing positive and negative markers for Cav1/CD44 vacuoles has moved to Supplemental Fig. S3B (formerly Supplemental Figure S1). Fonts and image sizes of figures 1-4 were corrected to ensure consistency and accuracy

