## Supplemental Figures for "CLIC-dependent internalization of caveolin-1 to lysosomal vacuoles in response to osmotic regulation"

### Supplemental Figure Legends

**Supplemental Figure S1. Quantification of CDC42-dependent Cav1 endocytosis**

Quantification of CDC42-GFP in MDA-MB-231 cells transfected with WT or DN CDC42-GFP were treated with hypotonic or isotonic media for 60 minutes and then fixed and labeled for Cav1 and CD44. The number of Cav1 endocytic vacuoles per cell and CDC42-GFP fluorescence per cell were quantified (n>30 whole cell images from three independent experiments; ANOVA with Tukey's post-test showing comparison between CDC42-GFP WT and DN; ****$p < 0.0001$; Scale bar: 2µm). See also Figure 1D.

**Supplemental Figure S2. Representative images from acid-wash internalization assay**

A) Representative STED images of all indicated conditions from Figure 2A (Scale bar: 2µm). B) Confocal images of representative conditions from Figure 2B (Scale bar: 10µm; Inset scale bar: 2µm).

**Supplemental Figure S3. Hypotonic shock induced Cav1/CD44 vacuoles are positive for endogenous LAMP2, GFP-GPI, and GLUT1 and negative for MVB markers and Transferrin receptor.**

A) Representative STED images of endogenous Cav1, CD44 where indicated, and either GFP-GPI, Transferrin receptor or GLUT1 in MDA-MB-231 cells incubated in either isotonic media or in hypotonic media for 60 minutes. (Scale bar: 5µm; Inset scale bar: 2µm). B) Representative STED images of endogenous Cav1, CD44 and either LAMP2, CD63 or LBPA in MDA-MB-231 cells incubated in either isotonic or hypotonic media for 60 minutes (Scale bar: 5µm; Inset scale bar: 2µm). Insets show triple labelled lysosomal vacuoles with colors as indicated.

**Supplemental Figure S4. Representative STED images of Cav1 vacuoles in Cav1-WT-HA and Cav1-K*R-HA cells**

Representative STED images of MDA-MB-231 cells expressing Cav1-WT-HA or Cav1-KR-HA incubated under isotonic or hypotonic media for 60 minutes from Fig. 3B (Scale bar: 2µm).

**Supplemental Figure S5. Representative STED images of MDA-MB-231, PC3 cells and Cavin1-GFP expressing PC3 cells treated with osmotic shock**

A) Representative STED images of MDA-MB-231 and PC3 cells treated with isotonic, hypotonic and hypertonic shock at indicated timepoints from Fig. 4B (Scale bar: 2 µm). B) Representative STED images of PC3 cells and PC3 cells stably transfected with Cavin1-GFP treated with isotonic, hypotonic and hypertonic shock at 60 minutes from Fig. 4C (Scale bar: 2 µm).

**Supplemental Figure S6. Cavin1 dissociates from Cav1 upon hypotonic shock**

MDA-MB-231 Cav1 KO cells expressing Cav1-myc and PC3 cells transiently expressing Cavin1-GFP were incubated in isotonic media for 60 minutes or hypotonic media for the indicated times before fixation and Cav1 and cavin1 labels imaged by TIRF widefield imaging. (MDA-MB-231: anti-myc for Cav1 and anti-Cavin1; PC3: anti-Cav1 and GFP for cavin1-GFP). Quantification of the Cav1 and cavin1 area masks and area overlap between the masks was quantified (Quantified from whole cell images, n ≥ 30 cells per condition from three independent experiments; ANOVA with Dunnett post-test comparing each condition to isotonic 4°C control; *$p < 0.05$; **$p < 0.01$; ***$p < 0.001$; ****$p < 0.0001$; Scale bar: 2 µm).

**Supplemental Figure S7. Representative confocal images of NHS-SS-biotin uptake experiments**

Representative confocal images of endocytosed biotin labeled with Alexa568 conjugated streptavidin in MDA-MB-231, PC3 cells and Cavin1-GFP expressing PC3 cells treated with isotonic, hypotonic and hypertonic shock at 60 minutes from Fig. 4 D (Scale bar: 5 µm).

**Supplemental Figure S8. Cav1 vacuoles form in cells seeded on PA hydrogels of defined elastic moduli spanning a range of physiologically relevant stiffnesses**

MDA-MB-231 and PC3 cells were seeded for 24 h on 140 Pa, 400 Pa, 2700 Pa, and 6000 Pa fibronectin-conjugated PA hydrogels, fixed and labelled for Cav1 and CD44. Representative STED images from 3 independent experiments showing individual Cav1 and CD44 channels are shown from Fig. 4 E. (Scale bar: 5 µm).

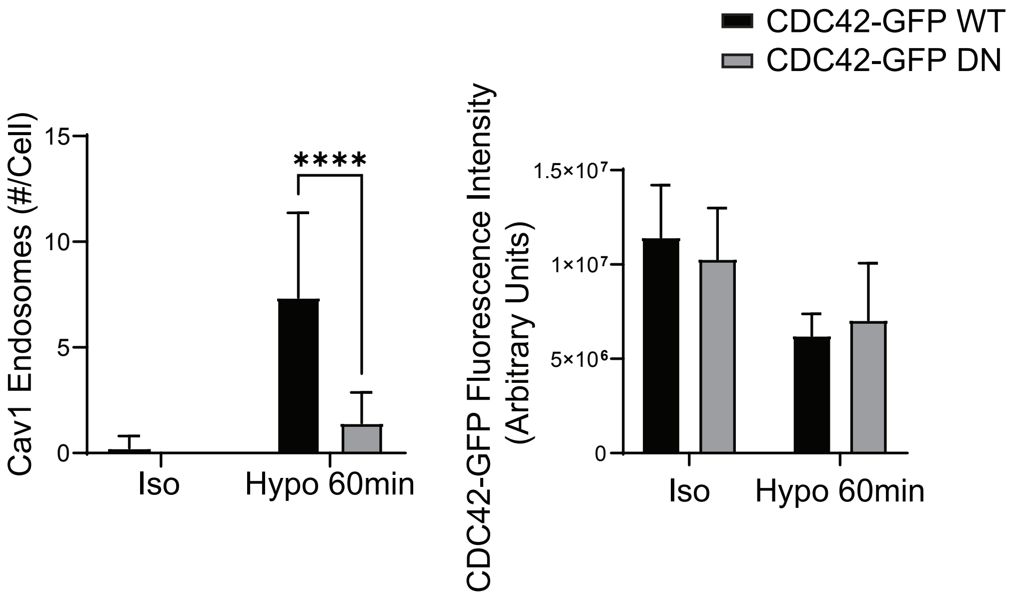


Wong et al Supp Fig S1

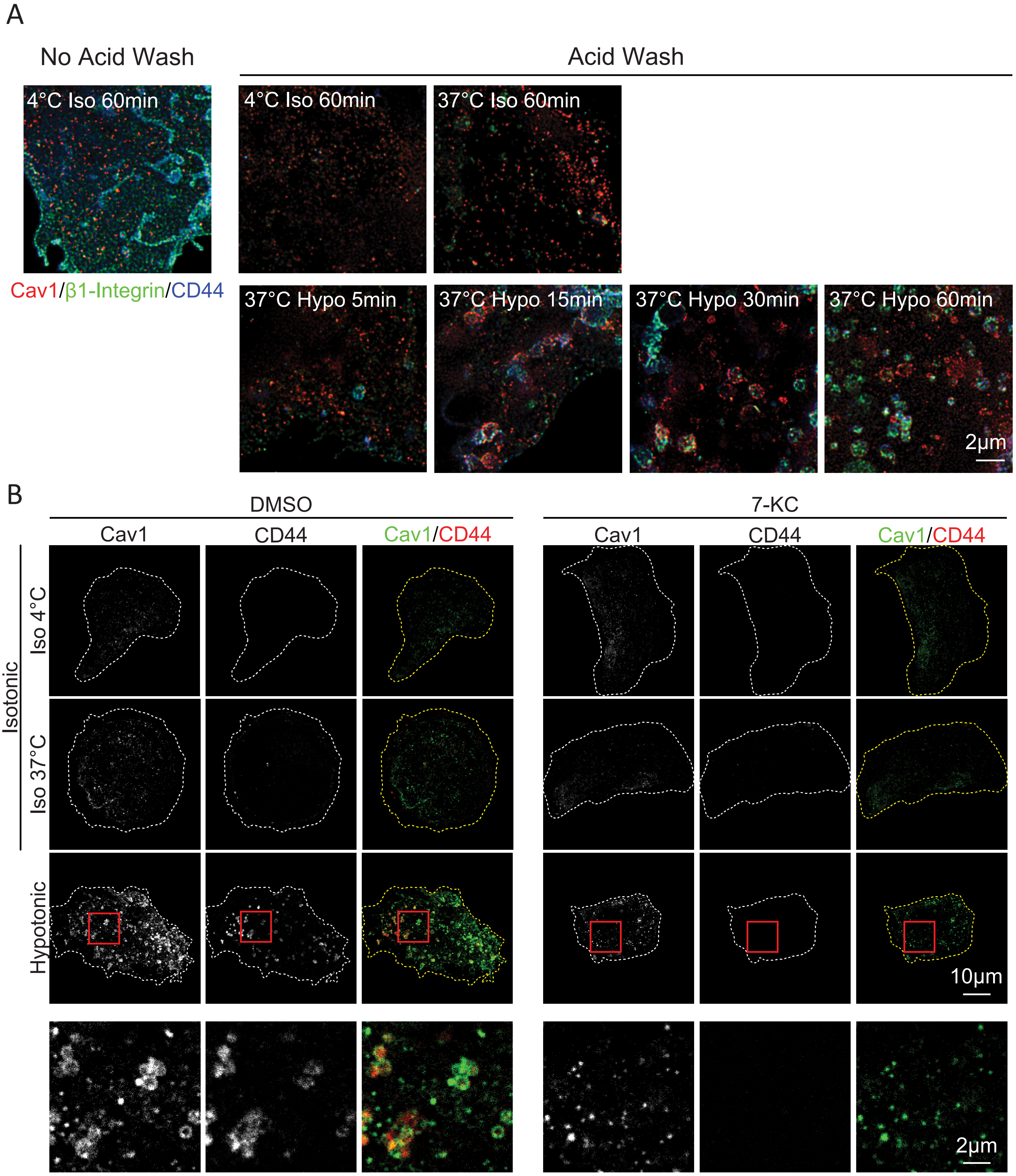


Wong et al Supp Fig S2

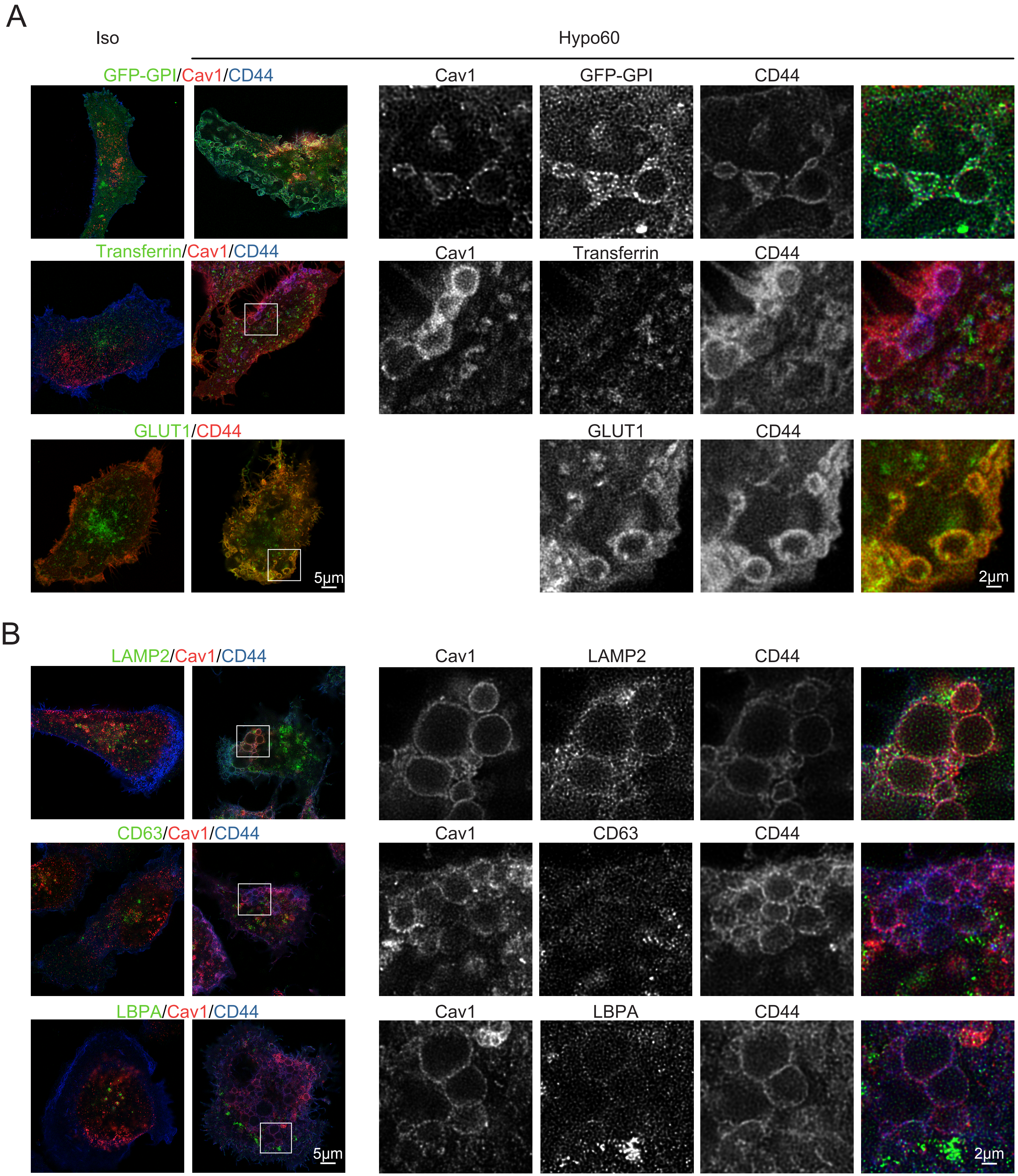


Wong et al Supp Fig S3

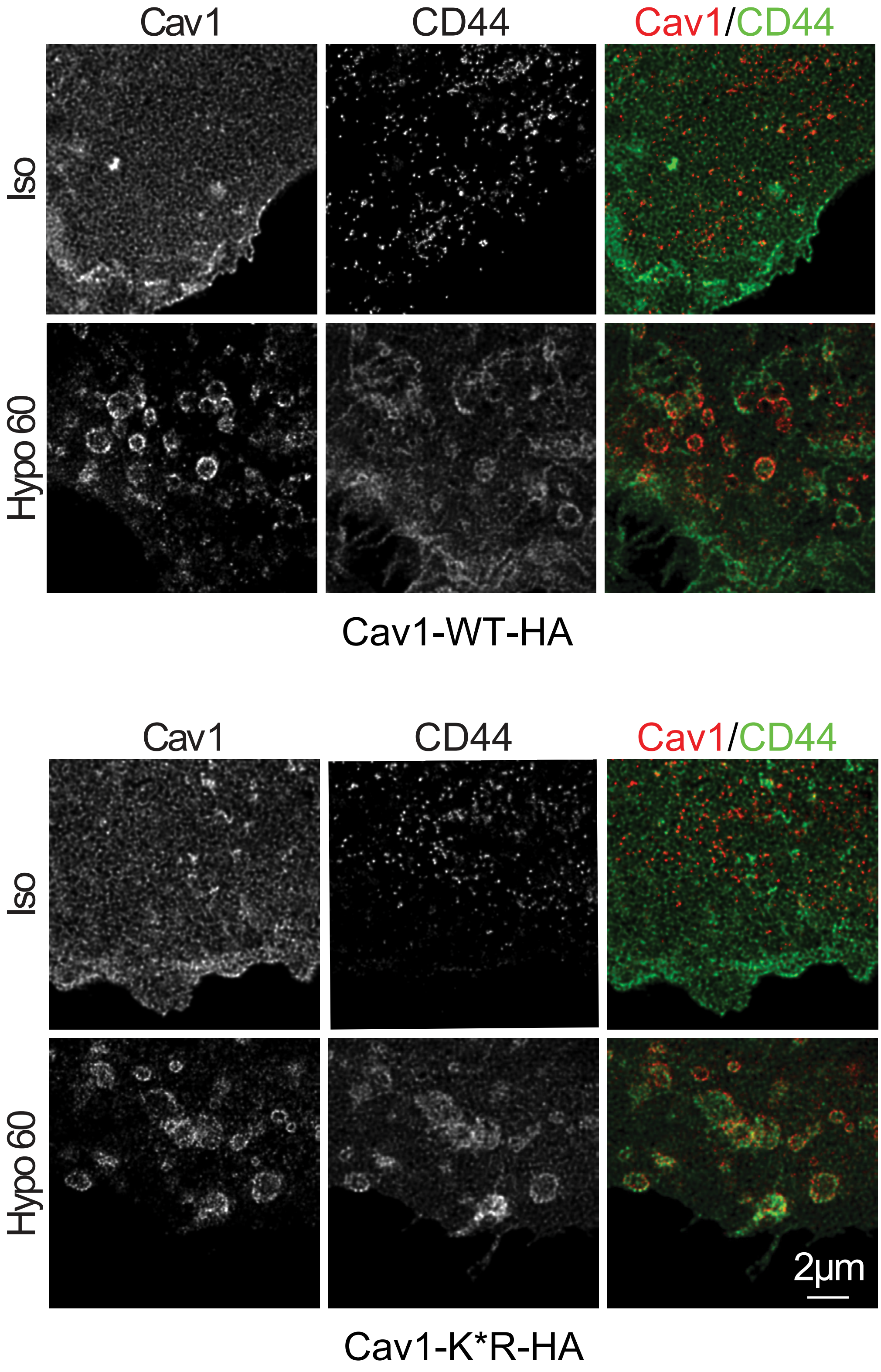


Wong et al Supp Fig S4

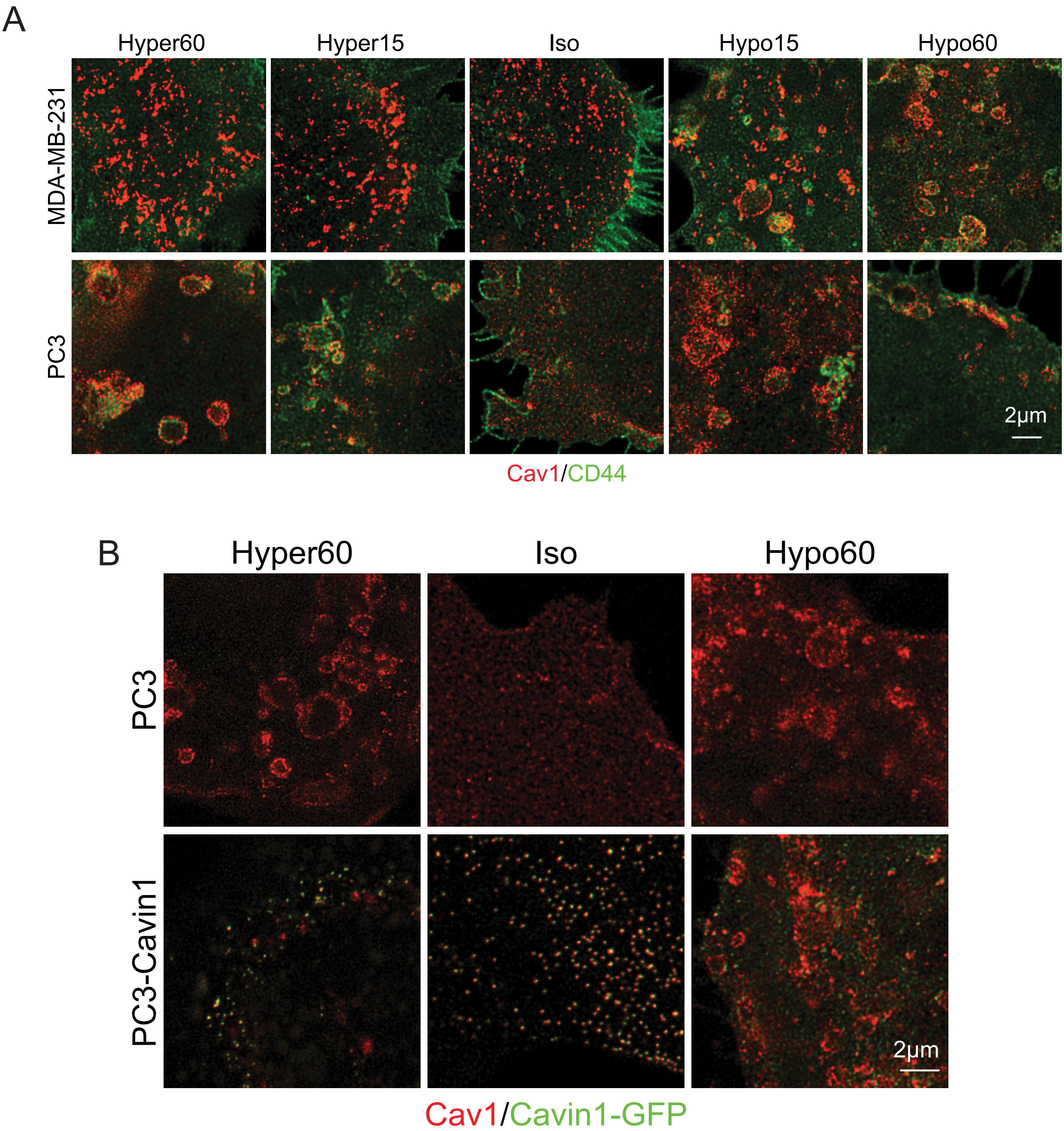


Wong et al Supp Fig S5

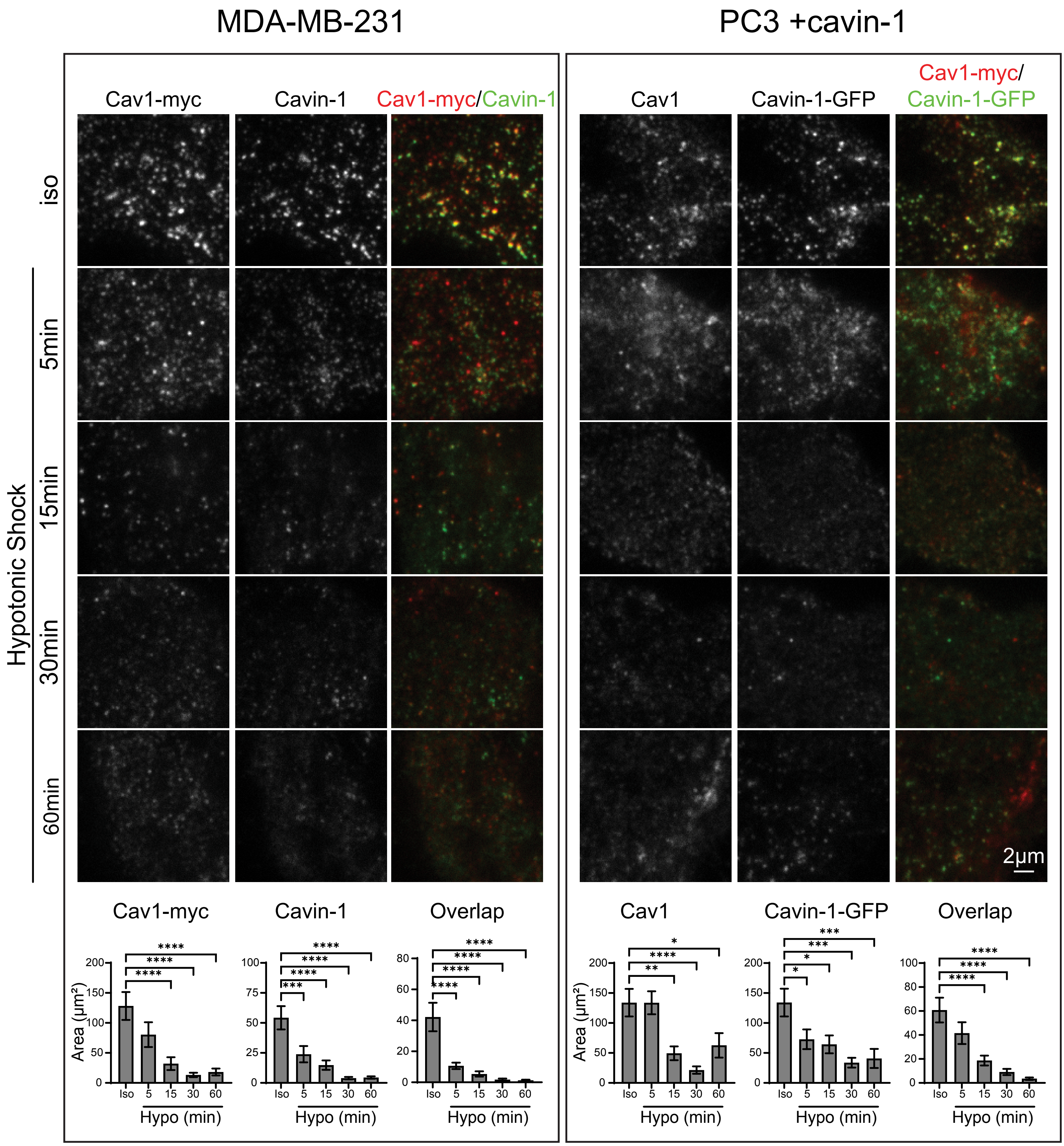


Wong et al Supp Fig S6

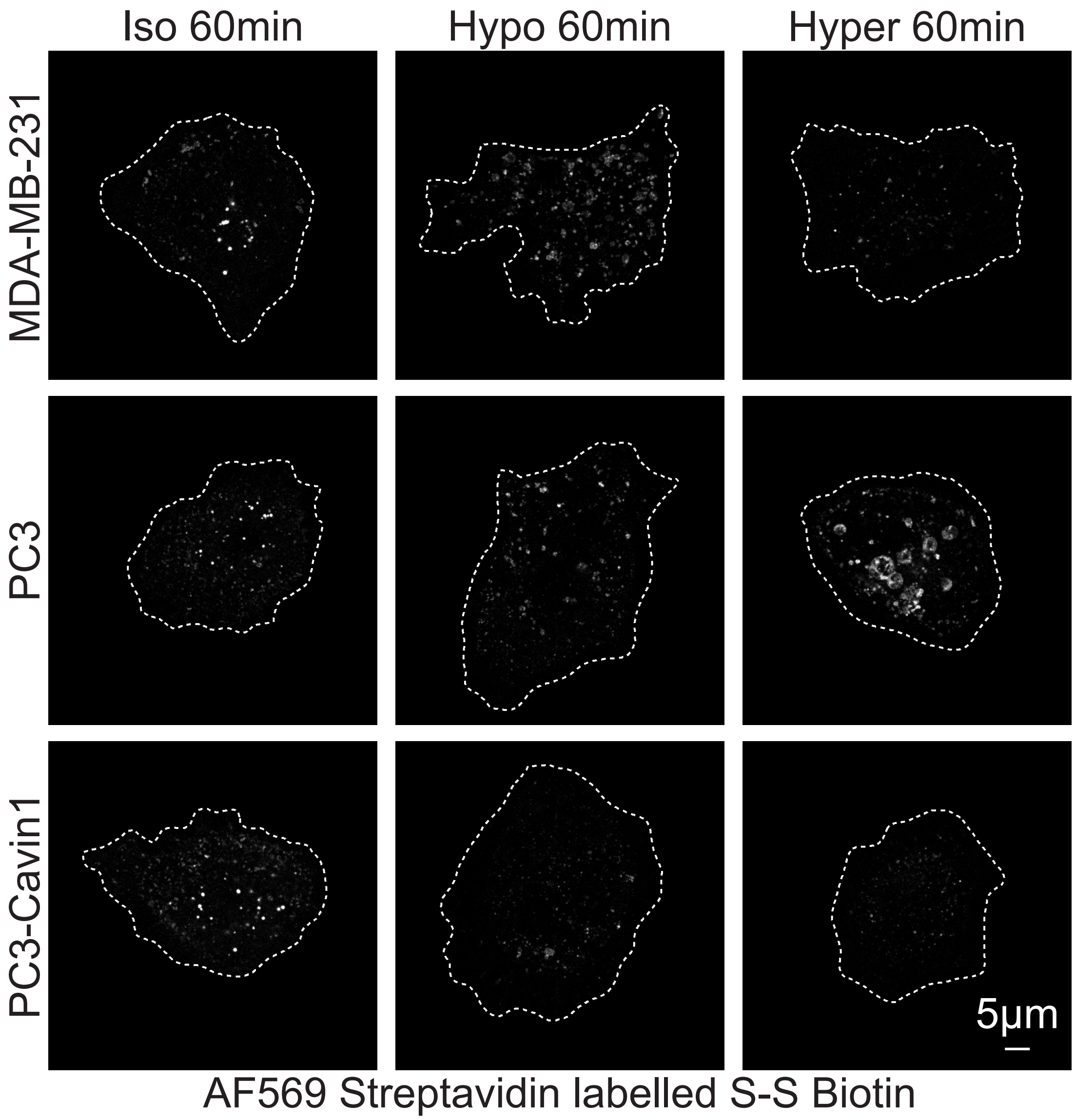


Wong et al Supp Fig S7

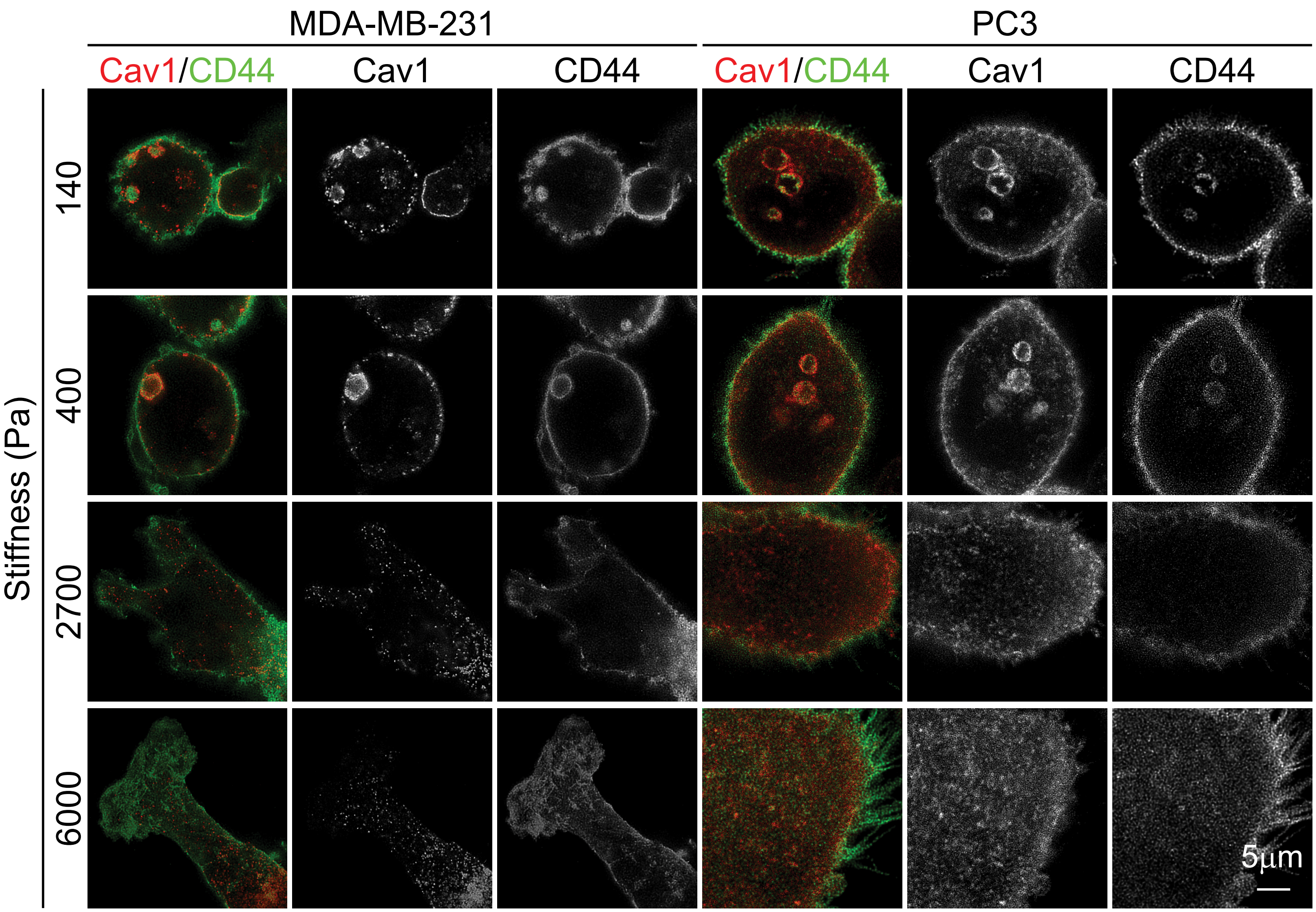


Wong et al Supp Fig S8
